# Land use and oriental pied hornbill occurrence in Singapore

**DOI:** 10.1101/2023.10.25.563926

**Authors:** Zaheedah Yahya, Min Yi Chin, Adlan Syaddad, Philip Johns

## Abstract

Oriental pied hornbills (*Anthracoceros albirostris)* disappeared from Singapore in the 1800s but were reintroduced in the 2000s. Since then, they have multiplied and even thrived in Singapore, spreading throughout much of the country. Because Singapore is extremely urban and densely populated, here we examine where hornbills have been seen as a function of land-use. We combine citizen science data from eBird and iNaturalist with GIS information to model hornbill sightings. We find that the percent Park and Beach cover have strong positive effects on the probability of hornbill sightings, and that percent Park cover has a positive interaction with percent Built-up and Sea cover. Our results are broadly consistent with sightings of oriental pied hornbills in the region.

## Introduction

Intense urbanization and deforestation have resulted in biodiversity loss (Corlett, 1992). Southeast Asia includes multiple biodiversity hotspots (De Bruyn et al., 2014) and also has one of the highest deforestation rates (Hughes, 2017). While birds are least dependent on Singapore’s nature reserves (Brook et al., 2003) some Singapore bird species have gone locally extinct. The oriental pied hornbill (*Anthracoceros albirostris)* disappeared from Singapore in the 1800s, and the comprehensive checklist published by Gibson-Hill in 1949 did not include these hornbills, even though this species survived in Peninsular Malaysia. In March 1994, after more than a century of absence, a pair was observed at Pulau Ubin, an island in Singapore’s northeast (Wong, 2011).

In 2006, Singapore’s National Parks Board (www.nparks.gov.sg) and the Wildlife Reserves of Singapore (now www.mandai.com/en/singapore-zoo.html) started the Singapore Hornbill Project to reintroduce the hornbills to Singapore (Cremades & Ng, 2012). The team set up artificial nests in previously unoccupied areas, such as southern Bukit Timah (Cremades et al., 2011). The program established populations in Bukit Timah, Pulau Ubin, and Singapore Botanic Garden within a few years (Cremades & Ng, 2012). Hornbills can survive fairly well in disturbed and fragmented habitats (Marsden & Pilgrim, 2003). Their successful adaptation to Singapore’s urban landscape is partly due to their use of cultivated fruit resources (Lok et. al., 2013; Tan, 2010).

Although the Singapore Hornbill Project was a success, the Singapore Red Data book lists the species as “Critically Endangered” (www.nparks.gov.sg/biodiversity/wildlife-in-singapore/species-list/bird) as of October 23 2023. Exploring the occupancy of the reintroduced hornbills, and relating hornbill sightings to land-use, has general implications to their reintroduction into urban environments.

Species distribution modelling (SDM) investigates relationships between species observations and environmental variables, assuming a functional relationship (Miller, 2010), and often using geographical information systems (GIS; Trisurat et al., 2013). Other studies of the oriental pied hornbills in Singapore focused on population distribution with sighting density maps (Strange & O’Dempsey, 2022). Citizen or Community Science platforms, such as eBird (Sullivan et al., 2009) and iNaturalist, can record observations of hornbill occupancy, especially in urban areas with high human density (Dell’amore, 2014). These platforms include metadata such as date, time and location, which allow the validation of observations through the citizen science community. Although there are potential biases, these databases can offer a viable alternative to scientific records (Tiago et al., 2017). Oriental pied hornbills are morphologically conspicuous and loud, and because high human population density correlates to sampling density (Prudic et al., 2017), hornbills in Singapore are a suitable object for citizen science.

Here, we explore land-use variables related to the presence of oriental pied hornbills. We hypothesize hornbills will thrive within grids with certain environmental features, and that these features should therefore be considered in conserving hornbills. We then compare these qualitative features to oriental pied hornbill sightings in the region.

## Materials and Methods

We utilized the citizen science platforms, iNaturalist (www.inaturalist.org/) and eBird (ebird.org), and downloaded records with spatial information from within Singapore. We excluded specific areas and data due to inaccessibility, such as islands, military restricted areas, and the ocean (Fig. 1a). Data from eBird (https://doi.org/10.15468/dl.pdf5eq) for oriental pied hornbill for the selected areas of Singapore, on August 31, 2022, yielded 5,195 approved observations from the eBird Basic Dataset from 1 January 2010 to 31 July 2022. We downloaded 1,089 research-grade sightings from iNaturalist using the query ‘species = Anthracoceros albirostris, location = Singapore’, from September 2011 to August 2022, from most parts of Singapore, on August 24, 2022 (Fig. 1b).

**Figure 1.**
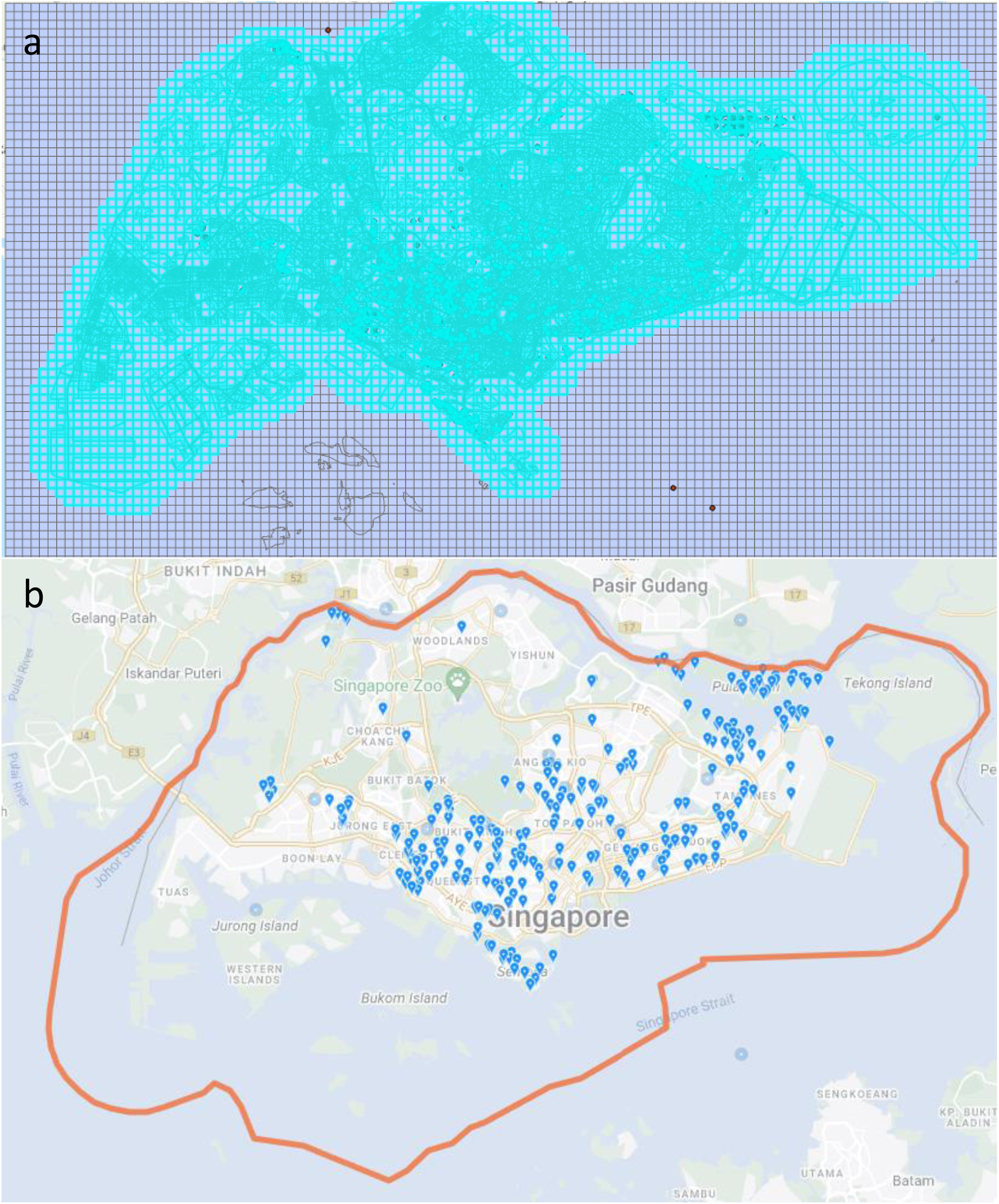
Map of selected study area. (a) Selected 0.5 km grids including Sentosa, excluding the Southern Islands and Bukom island. (b) iNaturalist map. Notice clusters at Pulau Ubin where hornbills were introduced.

We obtained geographical coordinates of sightings from the metadata of each sighting in the downloaded raw data and used those to map sightings on ArcGIS Pro 3.1.1 (www.esri.com). For land cover data, we included the Master Plan 2014 Land Use of Singapore, a vector polygon map extracted from Open Government Products (https://data.gov.sg), loaded as a base map. The map was published by Urban Redevelopment Authority (URA; https://www.ura.gov.sg/Corporate). The polygon map was indicative of each development land parcel with landscape characteristics and was last updated in 2017. The Coordinate Reference Systems used was the localized datum SVY21, commonly used in Singapore.

Four grid systems of various sizes of squares (0.5 to 2.5 km; Fig. A1) were generated as shapefiles and layered onto the Master Plan 2014 in ArcGIS. We obtained a list of spatial features in each grid by using the union tool in ArcGIS pro followed by the spatial join tool with the option *join one to one* to record sightings. The bounding area of the Master Plan map was defined by selecting the grids with the Select function on ArcGIS, followed with removal of grids made up entirely of sea, or inaccessible areas such as the military sites.

For each grid square we calculated the percentage land-use of the total area, as determined by Master Plan. The Master Plan 2014 Land Use of Singapore listed thirty-three zoning notations. Based on the zoning interpretations in The Planning Act Master Plan Written Statement 2019 (https://www.ura.gov.sg/-/media/Corporate/Planning/Master-Plan/MP19writtenstatement.pdf), we simplified this list into six broad land-use categories, adapted from the United States Geological Survey Level 1 Classification (Anderson et al. 1976): Sea, Waterbody, Agriculture, Built-up, Park and Beach (Table A1).

**Table 1.**
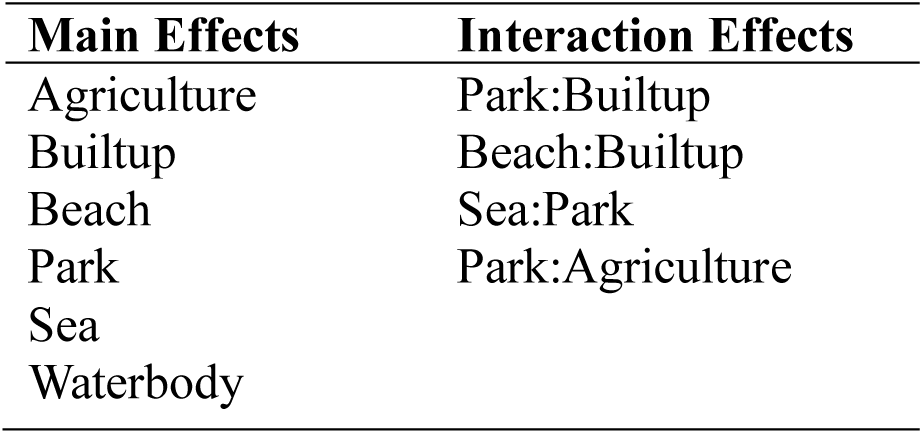
Initial main and interaction effects for GLM, before stepwise deletion of effects.

We estimated the correlation among these six categories using the R studio function *corrplot* (Wei & Simko, 2021) (Fig. B1).We constructed generalized linear models (GLM) for four grid sizes (0.5, 1.0, 2.0, and 2.5 km) with binomial error distribution (McCullagh & Nelder, 1989) in R Studio version 2023.06.0 (R Development Core Team, 2023), where sighting was a binary response variable; i.e., we did not include the number of sightings within a grid. We included the six land-use categories as main predictors and interaction terms between intercorrelated variables. We enhanced model performance with stepwise logistic regression (Kassambara, 2018) by iterative deletion of predictors based on statistical significance. We then ranked candidate models for each grid size by AIC and BIC (Burnham & Anderson, 2002). We predicted marginal effects of the selected model with other effects held constant at their mean values using the R package *sjPlot* with the function plot_model (Lüdecke, 2023).

## Results

We explored correlations among land uses for different grid sizes, 0.5 to 2.5 km. Correlation patterns were similar for all grid sizes, and we present the correlations among land uses for 2.5 km grids in which there were hornbill sightings (Fig. B1a) and all grids (Fig. B1b); patterns were similar for both (Fig. B1).

There were strong negative correlations between percent cover Sea and Built-up, between Park and Sea, and between Park and Built-up. There was a strong positive correlation between Beach and Sea, and between Park and Agriculture. We included these pairs of correlated land uses as possible interaction effects when modelling the probability of hornbill sighting (Table 1) before the stepwise elimination of terms (Table B1). We did not consider the strong correlations between Sea and Built-up, or Sea and Beach, as such correlations were to be expected irrespective of hornbill sightings.

Three of the four optimized GLM contained similar significant predictors and interaction terms (Table B2). For grid sizes 1.0 km, 2.0 km and 2.5 km, percent cover Beach, Sea and Park were significant predictors of hornbill sightings, in addition to the interaction terms of Built-up and Park, as well as Sea and Park.

**Table 2.** Summary statistics of percentage area of land use types of grids with sightings (*n* = 91) and without sightings (*n* = 114) for 2.5 km square grids.

| Land use type (%) | With Sighting |  |  |  |  |  | Without Sighting |  |  |  |  |  |
| --- | --- | --- | --- | --- | --- | --- | --- | --- | --- | --- | --- | --- |
|  | Mean | SD | SE | Median | Min | Max | Mean | SD | SE | Median | Min | Max |
| **Agriculture** | 1.05 | 4.90 | 0.51 | 0.00 | 0.00 | 32.51 | 0.53 | 3.88 | 0.36 | 0.00 | 0.00 | 40.15 |
| **Beach** | 0.12 | 0.35 | 0.04 | 0.00 | 0.00 | 1.81 | 0.01 | 0.08 | 0.01 | 0.00 | 0.00 | 0.85 |
| **Built-Up** | 65.41 | 31.07 | 3.26 | 78.89 | 0.53 | 99.45 | 32.83 | 34.73 | 3.25 | 19.35 | 0.00 | 100.00 |
| **Park** | 11.66 | 15.59 | 1.63 | 5.26 | 0.00 | 76.17 | 7.92 | 19.08 | 1.79 | 0.00 | 0.00 | 92.54 |
| **Waterbody** | 4.60 | 7.78 | 0.82 | 1.82 | 0.00 | 39.15 | 2.32 | 5.91 | 0.55 | 0.00 | 0.00 | 35.82 |
| **Sea** | 17.16 | 27.61 | 2.89 | 0.01 | 0.00 | 99.16 | 56.38 | 38.74 | 3.63 | 67.14 | 0.00 | 99.98 |

Comparing AIC and BIC (Burnham & Anderson, 2002) among the optimized models for all four grid sizes revealed the largest grid size (2.5 km) produced the lowest AIC (199.70) and BIC (219.63), whose values were approximately 90 and 92, respectively, less than the next best grid size (Table B3). We therefore consider the 2.5 km grid size for the rest of our analysis.

We summarized statistics of grids with and without sightings (Table 2). Hornbills were sighted in grids with a higher percentage of most land uses except Sea. Interestingly, hornbills were seen in grids with about twice as much Built-up (65.41%) and Waterbody (4.60%) land-use as grids without sightings (Table 2).

For the GLMs for 2.5 km grids, models 4.4 and 4.5 yielded the lowest AIC and BIC values; relative to the null model GLM 4.4 and 4.5 performed similarly (∂AIC = -82.94, -83.90 ; ∂BIC = -63.02, -67.30, respectively; Table B3). Model 4.4 included Waterbody, although the effect was not significant (p = 0.32; Table 3). The mean value of Waterbody percentage is about twice for grids with sightings vs without (Table 2). Because water presence influences bird diversity (Lerm et al, 2023), we retained Waterbody as an effect and consider GLM 4.4, which included four main predictors, Beach, Waterbody, Sea and Park, and two pairs of interacting terms, Built-up and Park, and Sea and Park (Table 3).

**Table 3.** GLM results of models 4.4 and 4.5, with 2.5 km grid size, for effects of Beach, Sea, Waterbody, Park and interaction effects of Park with Waterbody, and Sea with Park, on hornbill sighting probability

| Model | Explanatory Variables | AIC | BIC | Estimate | Standard Error | Pr(>\|z\|) |
| --- | --- | --- | --- | --- | --- | --- |
| **4.4** |  | 200.66 | 223.92 |  |  |  |
|  | Intercept |  |  | 0.35 | 0.34 | 0.30837 |
|  | Beach |  |  | 447.40 | 140.98 | 0.00151 ** |
|  | Sea |  |  | -3.58 | 0.72 | 7.88e-07 *** |
|  | Waterbody |  |  | 2.75<br>-4.76 | 2.77 | 0.3205 |
|  | Park |  |  | 16.92<br>14.90 | 1.84 | 0.00977 ** |
|  | Builtup:Park |  |  |  | 5.95 | 0.00445 ** |
|  | Sea:Park |  |  |  | 5.18 | 0.00404 ** |
| **4.5** |  | 199.70 | 219.63 |  |  |  |
|  | Intercept |  |  | 0.49 | 0.31 | 0.1183 |
|  | Beach |  |  | 446.83 | 141.03 | 0.0015 ** |
|  | Sea |  |  | -3.74 | 0.71 | 1.28e-07 *** |
|  | Park |  |  | -4.06<br>15.56 | 1.63 | 0.01294 * |
|  | Builtup:Park |  |  | 14.34 | 5.68 | 0.00618 ** |
|  | Sea:Park |  |  |  | 5.10 | 0.00494 ** |

Predicted marginal probabilities of hornbill sightings were negatively related to percent Sea in a grid (Fig. 2a), but positively related to percent Park, Beach and Waterbody (Fig. 2b-d). All else being equal, predicted probability of hornbill sighting rose from more than 25% in grids with no Park cover to more than 90% when Park cover was at least 50% (Fig. 2b). The strongest effect was due to percent Beach cover (Fig. 2c); very small increases in Beach resulted in large increases in marginal probability of sighting hornbills, although Beach never topped 1.81% in grids with sightings (or 0.85% without; Table 2).

**Figure 2.**
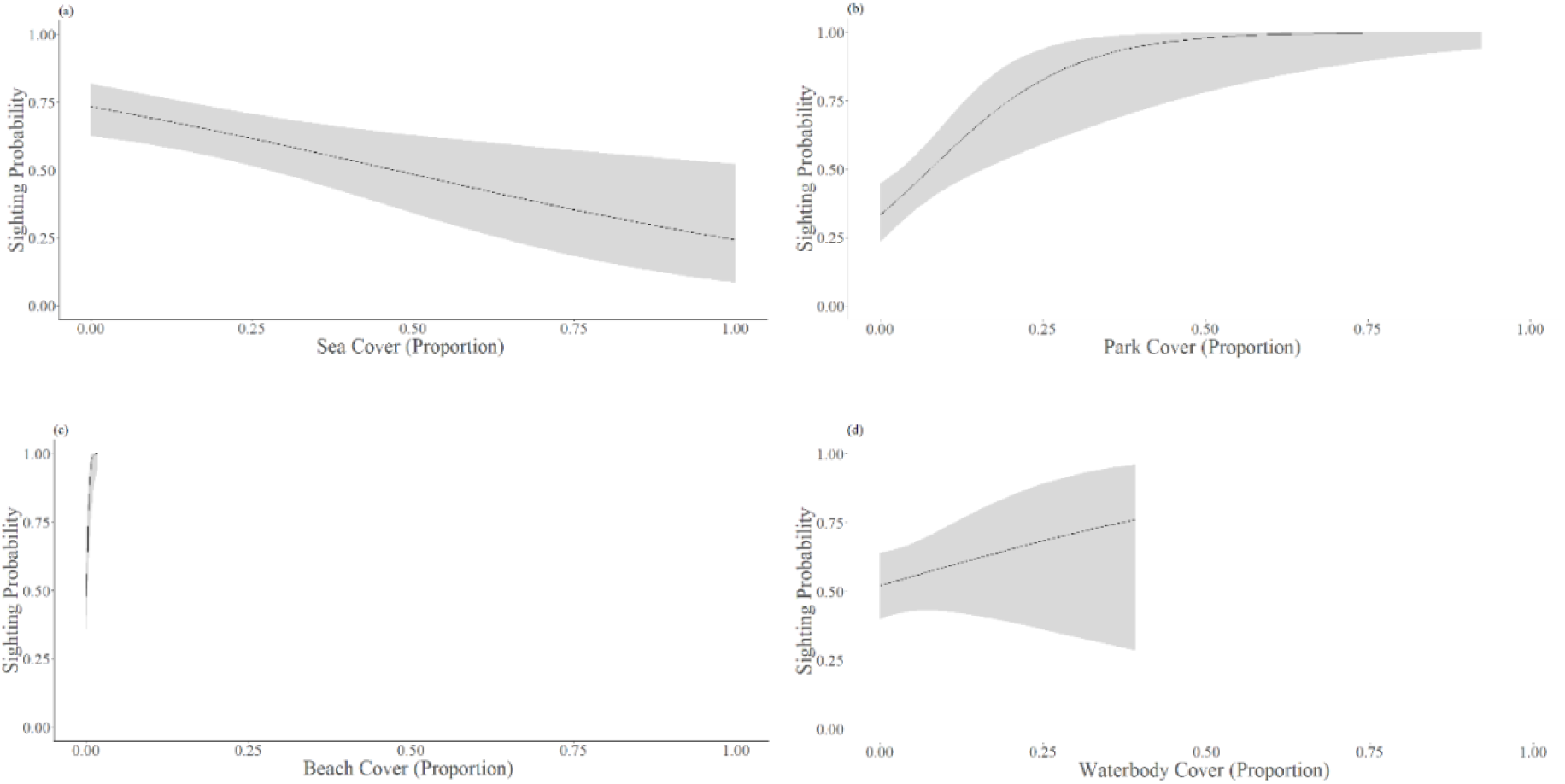
Marginal relationships between the predicted probability of hornbill sightings and land use for (a) Sea; (b) Park; (c) Beach; (d) Waterbody. Curves derived from GLM model 4.4 (Table 3); shaded regions show 95% confidence around each curve. X-axes constrained to observed values with hornbill sightings.

Predicted interactions between land-use types on the probability of hornbill sightings also reveal that Park is very important to the probability of hornbill sightings (Fig. 3). Grids with less Park show a stronger negative relationship between Sea and the probability of hornbill sighting (Fig. 3a). Similarly, the positive relationship between Built-up and hornbill sightings disappears as the percent Park decreases to zero (Fig. 3b).

**Figure 3.**
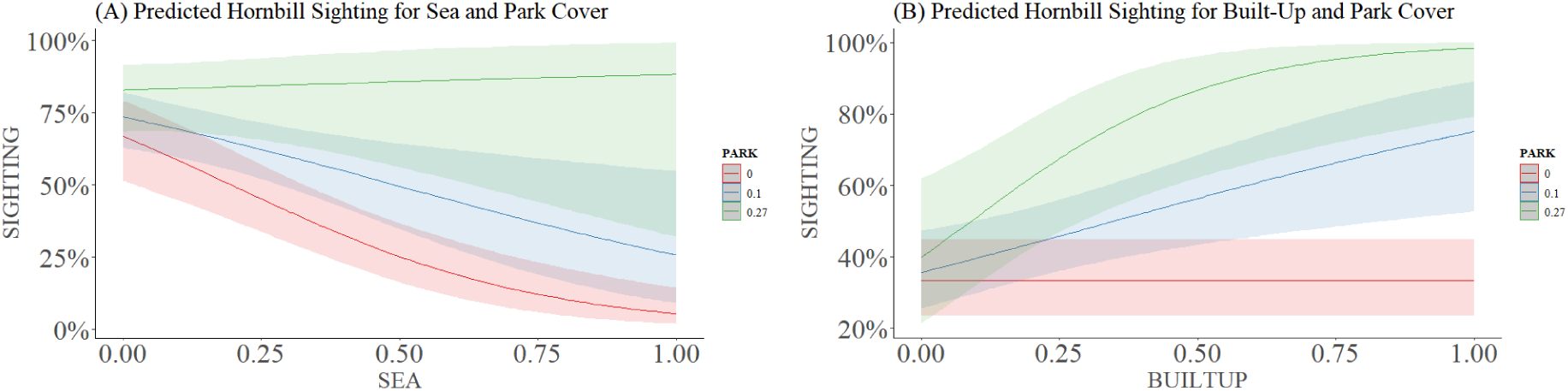
Predicted probabilities of hornbill sightings for the interactions effects between land use types: (a) Sea and Park; (b) Built-up area and Park. Curves show relationships predicted by GLM model 4.4 (Table 4); shaded areas show 95% confidence intervals around each curve. The average percent Park was 11.66% ± 15.59 SD (Table 2); curves for 0, 11, and 27% Park (Aiken et. al., 1991).

## Discussion

We used presence-only data to explore factors in land use that predict sighting of oriental pied hornbills in Singapore. Overall, the sighting of oriental pied hornbills in Singapore was influenced by Beach, Park, Waterbody and Sea.

These findings agree with published oriental pied hornbills’ habitat criteria (Kemp, 1995). Oriental pied hornbills tend to live at forest edges, coastal mangroves, and forested islands (Ng et al., 2011). Not surprisingly, hornbill sightings increased with percent Park; being a forest species, hornbills are found near forests. However, percent Park also influenced how other land uses affected hornbill sightings: Built-up was somewhat surprisingly positively related to hornbill sightings, but only if Park land was present on a grid.

Beach and Sea land use is also important for hornbill sighting, which underscores their habitat in the coastal areas. Beach is the strongest predictor with the largest effect. This positive relationship between hornbill sightings and beaches is probably partly due to hornbills living in coastal lowlands, and beaches often abut coasts. This pattern highlights the importance of urban land-use in conservation and management plans for hornbills; i.e., if hornbills live near beaches, then hornbill nest boxes near beaches might be particularly effective.

Park land-use includes non-forested areas in Singapore as well; e.g., Gardens by The Bay, West Coast Park, and East Coast Park. High human foot traffic at these popular parks might increase the chances of hornbill sightings. The percentage of Built-up area was twice a high in grids with sightings as in grids without sightings, and the percentage of Beach was ten times as high. These too might indicate a sampling bias favoring sightings where there are more people. So might the positive relationship between Built-up and Park; more observers in Built-up areas might be more likely to see hornbills. Alternatively, oriental pied hornbills seem to be very tolerant of human disturbance, and even perch on the rooves and balconies of housing estates. The positive interaction effect between Park and Built-up might genuinely reflect hornbill ecology. Observation bias has a larger influence in studies with small samples (Pearson et al. 2006). Our study included over 6,200 records, which should lessen bias.

Limiting the study to the presence of sightings, rather than using their frequency, should reduce the more egregious sampling biases. Despite potential biases, citizen science tools are useful, especially in places where large numbers of people have the opportunity to contribute observations (Lloyd et al., 2020; Hall et al., 2021), such as Singapore.

To validate the qualitative conclusions of our model, we explored the distribution of sightings of oriental pied hornbills on iNaturalist region-wide. We focused on three sites: Langkawi, Peninsular Malaysia; Miri, Eastern Malaysia; and Brunei. We used all observations as of October 05, 2023 (Fig. 4).

**Fig. 4.**
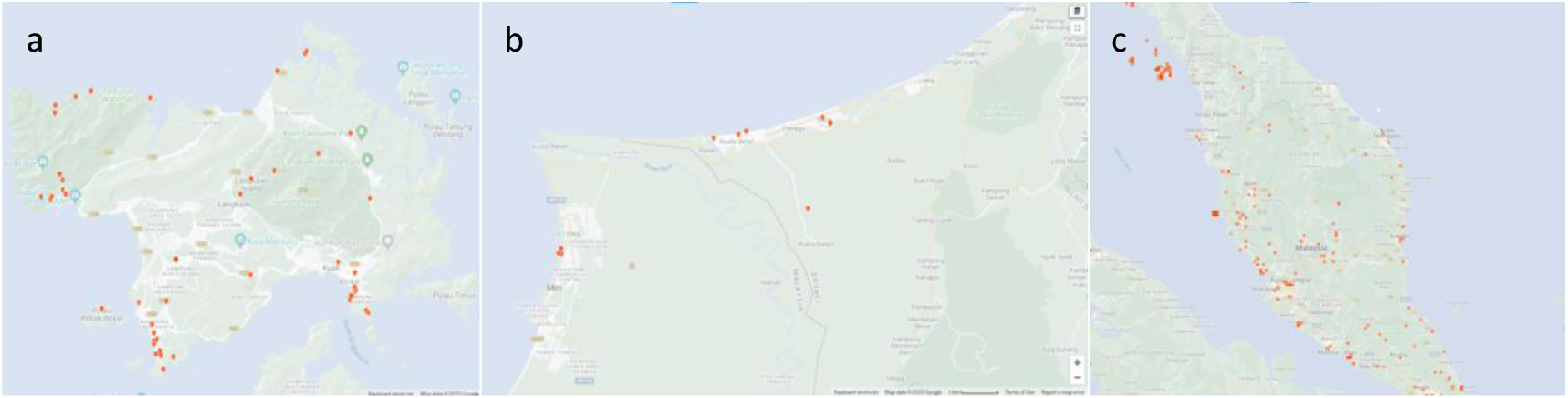
Maps of all records of oriental pied hornbill observations on iNaturalist in (a) Langkawi, Malaysia; (b) the west coast of Sarawak and Brunei; (c) Peninsular Malaysia. Red markings indicate research grade observations.

In Langkawi, sightings were abundant on the three beaches, along lowland coastal forests, and a few sightings (n = 4) were in the Wildlife Park in the center of the island (Fig. 4a). We found similar patterns in Miri, Malaysia, and in Brunei, where almost all sightings were near beaches (Fig. 4b). On a larger scale in Peninsular Malaysia, almost all sightings are in coastal regions and many are near beaches. Inland sightings are generally along waterways and on lower forests, typically less than 50 meters elevation (Fig. 4c).

These sightings (Fig. 4) agree with our analysis of Singapore. In general, we see hornbills in Singapore in locations consistent with the region, even though Singapore is extremely urban.

## Conclusion

The model we describe estimates land-use that predicts the probability of oriental pied hornbills sighting. Predictors are largely consistent with where hornbills are found outside of cities, although there are positive interactions between Built-up and Park cover on hornbill sightings. Our analysis demonstrates that citizen science data may be valuable in predicting species sighting. Conservation managers can use similar analyses for predicting land-use important to other urban wildlife. Our study will also assist conservation managers in land-scarce Singapore where there are many competing land uses.

## Appendix A: Methodological additions

**Fig. A1.**
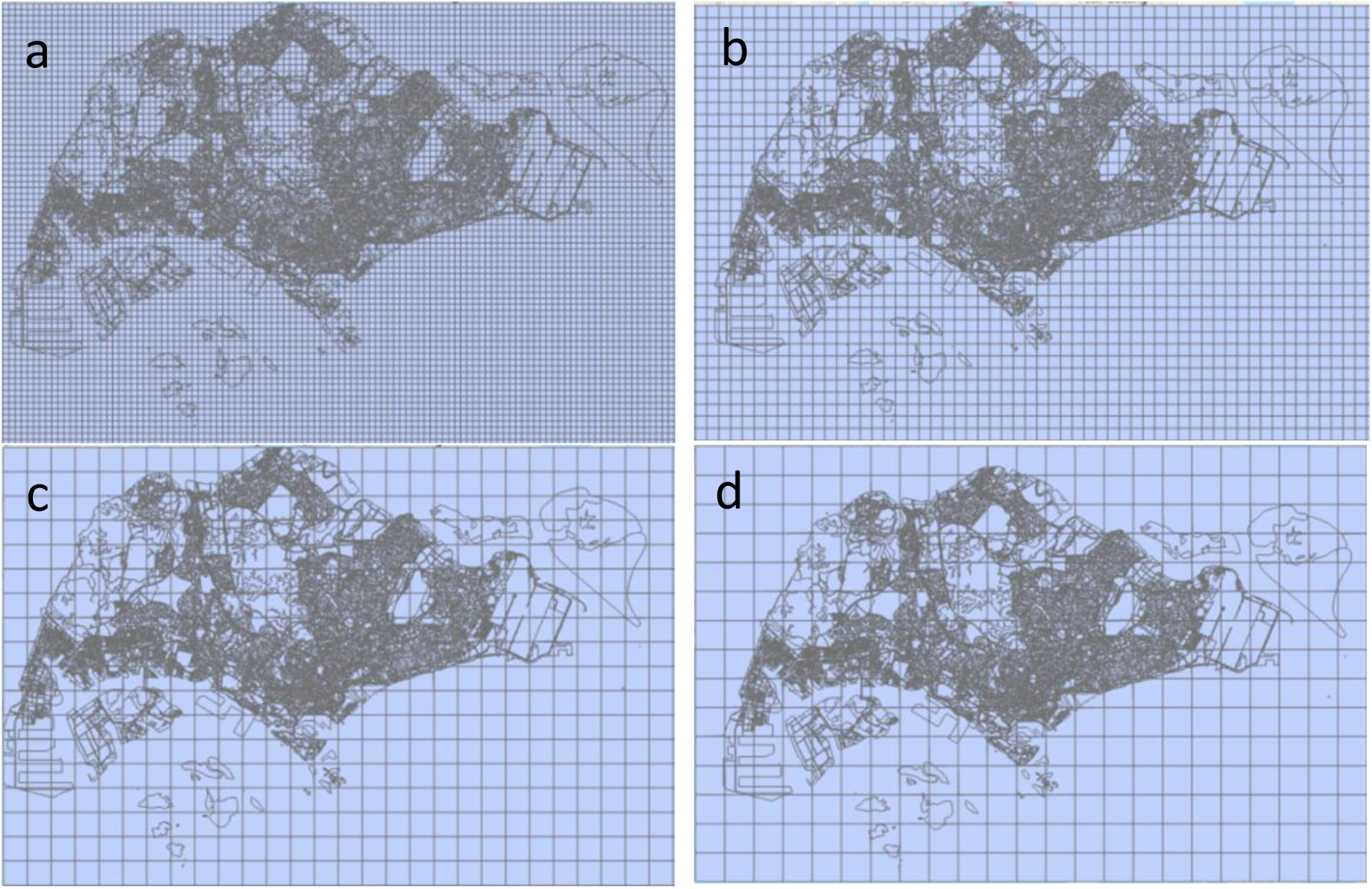
Map of study area with four grid sizes used in this study: (a) 0.5 km (b) 1.0 km (c) 2.0 km (d) 2.5 km on a side.

**Table A1.** Potential Predictors for hornbill sighting probability in Master Plan 2014 Land Use of Singapore.

| Zoning | General Category |
| --- | --- |
| Agriculture | Agriculture |
| Beach | Beach |
| Business 1 | Built-Up |
| Business 1 – White<sup>1</sup> | Built-Up |
| Business 2 | Built-Up |
| Business 2 - White | Built-Up |
| Business Park | Built-Up |
| Business Park - White | Built-Up |
| Cemetery | Built-Up |
| Civic & Community Institution | Built-Up |
| Commercial | Built-Up |
| Commercial & Residential | Built-Up |
| Educational Institution | Built-Up |
| Health & Medical Care | Built-Up |
| Hotel | Built-Up |
| Light Rapid Transit | Built-Up |
| Mass Rapid Transit | Built-Up |
| Place Of Worship | Built-Up |
| Port / Airport | Built-Up |
| Reserve Site | Built-Up |
| Residential | Built-Up |
| Residential / Institution | Built-Up |
| Residential With Commercial At 1St Storey | Built-Up |
| Road | Built-Up |
| Special Use<sup>2</sup> | Built-Up |
| Sports & Recreation | Built-Up |
| Transport Facilities | Built-Up |
| Utility | Built-Up |
| White | Built-Up |
| Open Space | Park |
| Park | Park |
| Waterbody | Waterbody |
| Sea | Sea |

<sup>1</sup>White zone denoted as a mixed use development
<sup>2</sup>Special Use denoted as no access zone for military purposes

## Appendix B. Results additions

**Fig. B1.**
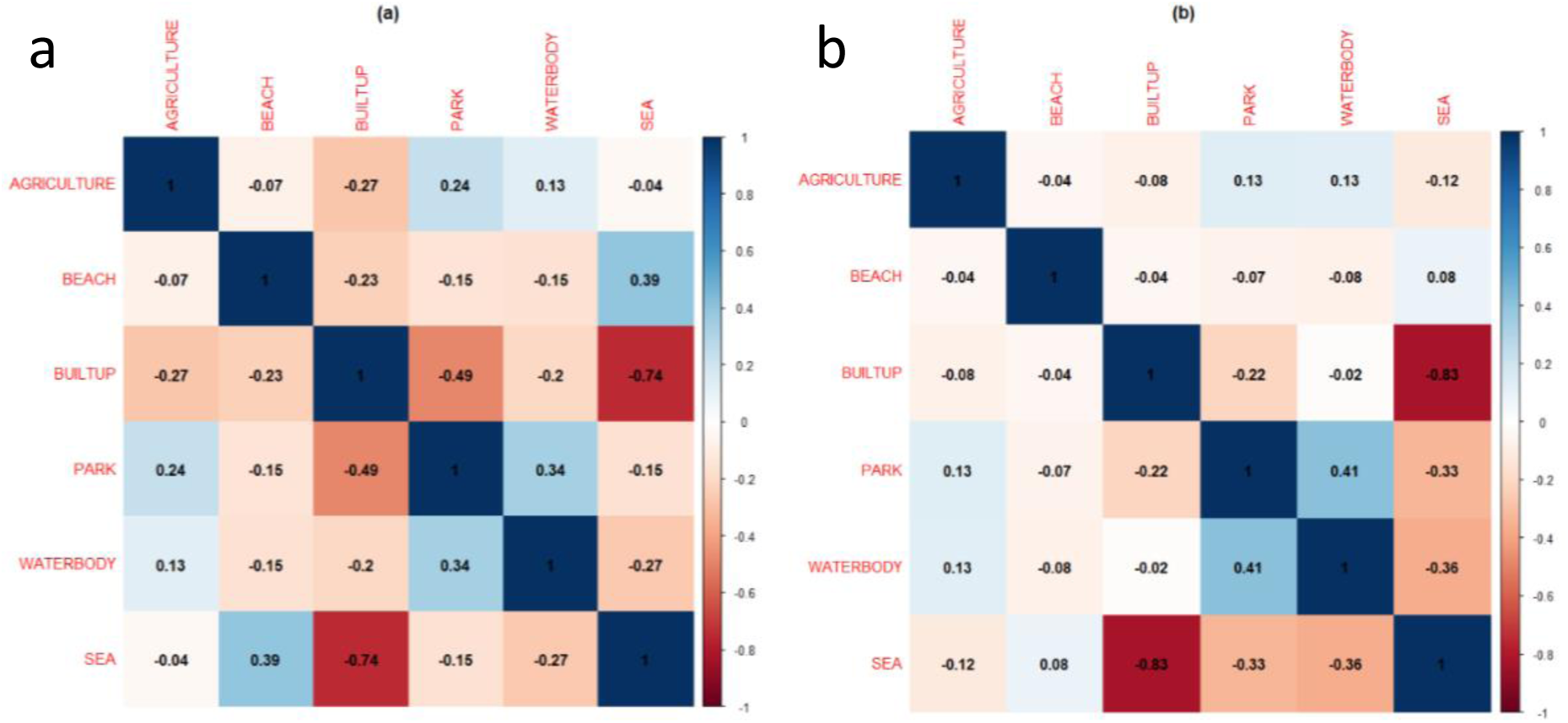
Correlation heatmap for all land use variables for 2.5 km grids: (a) with hornbill sightings only, (b) for all grids. Blue and red denote positive and negative correlations, respectively; correlation coefficient (r) included for each pair of land use variables.

**Table B1.** Stepwise deletion of non-significant effects in GLM models of hornbill sighting probability for grid squares of sizes 0.5 km, 1.0 km, 2.0 km, 2.5 km.

| Grid Size | 1<sup>st</sup> Iteration | 2<sup>nd</sup> Iteration | 3<sup>rd</sup> Iteration | 4<sup>th</sup> Iteration | 5<sup>th</sup> Iteration | 6<sup>th</sup> Iteration |
| --- | --- | --- | --- | --- | --- | --- |
| 0.5 km | Beach:Builtup | Agriculture | Sea |  |  |  |
| 1.0 km | Beach:Builtup | Builtup | Waterbody | Agriculture |  |  |
| 2.0 km | Beach:Builtup | Park:Agriculture | Agriculture | Builtup | Waterbody | Beach |
| 2.5 km | Beach:Builtup | Park:Agriculture | Agriculture | Builtup | Waterbody |  |

**Table B2.** Optimized final models for probability in hornbill sighting as response variable for the grid scale of 0.5 km, 1.0 km, 2.0 km, and 2.5 km squares with predictor variables.

| Scale (km) | Explanatory Variables | AIC | BIC | Estimate | Standard Error | Pr(>\|z\|) |
| --- | --- | --- | --- | --- | --- | --- |
| **0.5** |  | 2555.52 | 2604.66 |  |  |  |
|  | Intercept |  |  | -3.97 | 0.32 | < 2e-16 *** |
|  | Beach |  |  | 39.88 | 6.59 | 1.44e-09 *** |
|  | Builtup |  |  | 2.11 | 0.34 | 3.89e-10 *** |
|  | Waterbody |  |  | 1.18 | 0.55 | 0.032 * |
|  | Park |  |  | 1.46 | 0.39 | 0.00017 *** |
|  | Builtup:Park |  |  | 5.78 | 0.90 | 1.21e-10 *** |
|  | Sea:Park |  |  | 10.79 | 1.86 | 6.17e-09 *** |
|  | Park:Agriculture |  |  | 18.46 | 3.10 | 2.65e-09 *** |
| **1.0** |  | 955.14 | 989.13 |  |  |  |
|  | Intercept |  |  | -0.93 | 0.11 | < 2e-16 *** |
|  | Beach |  |  | 201.74 | 44.33 | 2.33e-06 *** |
|  | Sea |  |  | -2.97 | 0.49 | 1.56e-9 *** |
|  | Park |  |  | -1.91 | 0.50 | 0.00012 *** |
|  | Builtup:Park |  |  | 7.74 | 1.56 | 7.15e-07 *** |
|  | Sea:Park |  |  | 14.79 | 3.47 | 2.05e-05 *** |
|  | Park:Agriculture |  |  | 21.89 | 6.98 | 0.00171 ** |
| **2.0** |  | 289.73 | 311.12 |  |  |  |
|  | Intercept |  |  | -0.08 | 0.23 | 0.708855 |
|  | Beach |  |  | 1613.06 | 1585.19 | 0.308879 |
|  | Sea |  |  | -2.79 | 0.62 | 8.94e-05 *** |
|  | Park |  |  | -4.64 | 1.42 | 0.000553*** |
|  | Builtup:Park |  |  | 13.09 | 4.00 | 0.0.00041 *** |
|  | Sea:Park |  |  | 21.59 | 6.06 | 0.000147 *** |
| **2.5** |  | 199.70 | 219.63 |  |  |  |
|  | Intercept |  |  | 0.49 | 0.31 | 0.1183 |
|  | Beach |  |  | 446.83 | 141.03 | 0.0015 ** |
|  | Sea |  |  | -3.74 | 0.71 | 1.28e-07 *** |
|  | Park |  |  | -4.06 | 1.63 | 0.01294 * |
|  | Builtup:Park |  |  | 15.56 | 5.68 | 0.00618 ** |
|  | Sea:Park |  |  | 14.34 | 5.10 | 0.00494 ** |

**Table B3.** Stepwise elimination of GLM models of hornbill sighting probability for grid squares of sizes 0.5 km, 1.0 km, 2.0 km, 2.5 km.

| Model<br>(Grid<br>Size) | Explanatory Variables | Number of<br>parameters | AIC | $\partial$AIC | BIC | $\partial$BIC |
| --- | --- | --- | --- | --- | --- | --- |
| **Null**<br>**Model**<br>**(0.5 km)** | No variables | 0 parameters | 2730.01 | 0.00 | 2736.19 | 0.00 |
| **1**<br>**(0.5 km)** | Agriculture + Beach + Builtup<br>+ Sea + Waterbody + Park +<br>Builtup*Park + Sea*Park +<br>Beach*Builtup +<br>Park*Agriculture | 6 main<br>parameters,<br>4<br>interactions | 2556.54 | -173.47 | 2624.10 | -112.09 |
| **1.1**<br>**(0.5 km)** | Agriculture + Beach + Builtup + Sea + Waterbody + Park + Builtup*Park + Sea*Park + Park*Agriculture | 6 main parameters, 3 interactions | 2556.79 | -173.22 | 2618.21 | -117.98 |
| **1.2**<br>**(0.5 km)** | Beach + Builtup + Sea + Waterbody + Park + Builtup*Park + Sea*Park + Park*Agriculture | 5 main parameters, 3 interactions | 2557.52 | -172.49 | 2612.80 | -123.39 |
| **1.3**<br>**(0.5 km)** | Beach + Builtup + Waterbody + Park + Builtup*Park + Sea*Park + Park*Agriculture | 4 main parameters, 3 interactions | 2555.52 | -174.49 | 2604.66 | -131.53 |
| **Null Model**<br>**(1.0 km)** | No variables | 0 parameters | 1094.36 | 0.00 | 1099.22 | 0.00 |
| **2**<br>**(1.0 km)** | Agriculture + Beach + Builtup + Sea + Waterbody + Park + Builtup*Park + Sea*Park + Beach*Builtup + Park*Agriculture | 6 main parameters, 4 interactions | 953.75 | -140.61 | 1007.16 | -92.06 |
| **2.1**<br>**(1.0 km)** | Agriculture + Beach + Builtup + Sea + Waterbody + Park + Builtup*Park + Sea*Park + Park*Agriculture | 6 main parameters, 3 interactions | 953.37 | -140.99 | 1001.93 | -97.29 |
| **2.2**<br>**(1.0 km)** | Agriculture + Beach + Sea + Waterbody + Park + Builtup*Park + Sea*Park + Park*Agriculture | 5 main parameters, 3 interactions | 956.88 | -137.48 | 1000.57 | -98.65 |
| **2.3**<br>**(1.0 km)** | Agriculture + Beach + Sea + Park + Builtup*Park + Sea*Park + Park*Agriculture | 4 main parameters, 3 interactions | 955.59 | -138.77 | 994.44 | -104.78 |
| **2.4**<br>**(1.0 km)** | Beach + Sea + Park + Builtup*Park + Sea*Park + Park*Agriculture | 3 main parameters, 3 interactions | 955.14 | -139.22 | 989.13 | -110.09 |
| **Null Model**<br>**(2.0 km)** | No variables | 0 parameters | 357.36 | 0.00 | 360.92 | 0.00 |
| **3**<br>**(2.0 km)** | Agriculture + Beach + Builtup + Sea + Waterbody + Park + Builtup*Park + Sea*Park + Beach*Builtup + Park*Agriculture | 6 main parameters, 4 interactions | 290.91 | -66.45 | 330.12 | -30.80 |
| **3.1**<br>**(2.0 km)** | Agriculture + Beach + Builtup + Sea + Waterbody + Park + Builtup*Park + Sea*Park + Park*Agriculture | 6 main parameters, 3 interactions | 289.08 | -59.28 | 324.73 | -36.19 |
| **3.2**<br>**(2.0 km)** | Agriculture + Beach + Builtup + Sea + Waterbody + Park + Builtup*Park + Sea*Park | 6 main parameters, 2 interactions | 294.22 | -63.14 | 326.30 | -34.62 |
| **3.3**<br>**(2.0 km)** | Beach + Builtup + Sea<br>+Waterbody + Park +<br>Builtup*Park + Sea*Park | 5 main<br>parameters,<br>2<br>interactions | 292.39 | -64.97 | 320.91 | -40.01 |
| **3.4**<br>**(2.0 km)** | Beach + Sea +Waterbody +<br>Park + Builtup*Park +<br>Sea*Park | 4 main<br>parameters,<br>2<br>interactions | 291.27 | -66.09 | 316.22 | -44.70 |
| **3.5**<br>**(2.0 km)** | Beach + Sea + Park +<br>Builtup*Park + Sea*Park | 3 main<br>parameters,<br>2<br>interactions | 289.73 | -67.63 | 311.12 | -49.80 |
| **Null**<br>**model**<br>**(2.5 km)** | No variables | 0 parameters | 283.60 | 0.00 | 286.93 | 0.00 |
| **4**<br>**(2.5 km)** | Agriculture + Beach + Builtup<br>+ Sea + Waterbody + Park +<br>Builtup*Park + Sea*Park +<br>Beach*Builtup +<br>Park*Waterbody | 6 main<br>parameters,<br>4<br>interactions | 203.89 | -79.71 | 240.44 | -46.49 |
| **4.1**<br>**(2.5 km)** | Agriculture + Beach + Builtup<br>+ Sea +Waterbody + Park +<br>Builtup*Park +<br>Park*Waterbody | 6 main<br>parameters,<br>3<br>interactions | 202.99 | -80.61 | 236.22 | -50.71 |
| **4.2**<br>**(2.5 km)** | Agriculture + Beach + Builtup<br>+ Sea +Waterbody + Park +<br>Builtup*Park + Sea*Park | 6 main<br>parameters,<br>2<br>interactions | 204.61 | -78.99 | 234.51 | -52.42 |
| **4.3**<br>**(2.5 km)** | Beach + Builtup + Sea<br>+Waterbody + Park +<br>Builtup*Park + Sea*Park | 5 main<br>parameters,<br>2<br>interactions | 202.66 | -80.94 | 229.25 | -57.68 |
| **4.4**<br>**(2.5 km)** | Beach + Sea +Waterbody +<br>Park + Builtup*Park +<br>Sea*Park | 4 main<br>parameters,<br>2<br>interactions | 200.66 | -82.94 | 223.92 | -63.02 |
| **4.5**<br>**(2.5 km)** | Beach + Sea + Park +<br>Builtup*Park + Sea*Park | 3 main<br>parameters,<br>2<br>interactions | 199.70 | -83.90 | 219.63 | -67.30 |

## Acknowledgements

We would like to thank the citizen scientists of Singapore who have contributed to the eBird and iNaturalist databases. Portions of this study were conducted by Zaheedah in partial fulfilment of MSC requirements for BCNBCS, DBS, NUS. Ms Michelle Quak Song Yun from NUS Libraries has been very helpful in assisting with GIS, Dr Ian Zhi Wen Chan from Department of Biological Sciences, NUS for assistance on data analysis, and Ms Bee Choo Strange from Hornbill Research Foundation who provided insights to this study. The work was partly supported in part by the Singapore Ministry of Education through the Yale-NUS College grant R-607-265-226-121, and by Yale-NUS College Centre for International and Professional Experience (CIPE).

## Ethical approval

This study was purely observational; animals were not handled, manipulated, or fed in any way. Many of the observations were carried out by community scientists in Singapore.

## References

Aiken, L. S., West, S. G., & Reno, R. R. (1991). Multiple regression: Testing and interpreting interactions. SAGE.

Anderson, J.R., Hardy, E.E., Roach, J.T., & Witmer, R.E. (1976). A land use and land cover classification system for use with remote sensor data.

Brook, B. W., Sodhi, N. S., & Ng, P. K. (2003). Catastrophic extinctions follow deforestation in Singapore. Nature, 424(6947), 420–423. 10.1038/nature01795

Burnham, K.P. & Anderson, D.R., 2002. Model selection and multimodel inference: a practical information-theoretic approach. *Springer Science & Business Media*, New York

Corlett, R. T. (1992). The ecological transformation of Singapore, 1819-1990. Journal of Biogeography, 19(4), 411. 10.2307/2845569

Cremades, M., Wong, H. L. T., Koh, S., Segran, R., Ng, S. (2011) Re-Introduction Of The Oriental Pied Hornbill In Singapore, With Emphasis On Artificial Nests. The Raffles Bulletin Of Zoology. Supplement No. 24: 5–10

Cremades, M. & Ng, S. C. (2012). Hornbills in the city: A conservation approach to hornbill study in Singapore. National Parks Board Singapore.

De Bruyn, M., Stelbrink, B., Morley, R. J., Hall, R., Carvalho, G. R., Cannon, C. H., Van den Bergh, G., Meijaard, E., Metcalfe, I., Boitani, L., Maiorano, L., Shoup, R., & Von Rintelen, T. (2014). Borneo and Indochina are major evolutionary hotspots for Southeast Asian biodiversity. Systematic Biology, 63(6), 879–901. 10.1093/sysbio/syu047

eBird Basic Dataset. Version: EBD_relJul-2022. Cornell Lab of Ornithology, Ithaca, New York. Jul 2022. 10.15468/dl.pdf5eq

Gibson-Hill, C. A. (1950). A checklist of birds of Singapore island.

Hall, M. J., Martin, J. M., Burns, A. L., & Hochuli, D. F. (2021). Ecological insights into a charismatic bird using different citizen science approaches. Austral Ecology, 46(8), 1255–1265. 10.1111/aec.13062

Hughes, A. C. (2017). Understanding the drivers of Southeast Asian biodiversity loss. Ecosphere, 8(1):e01624.10/1002/ecs2.1624

iNaturalist community. Observations of Anthracoceros albirostris from Singapore, observed between September 2011 to August 2022. Exported from https://www.inaturalist.org on August 24, 2022.

Kassambara, A. (2018). Machine learning essentials: Practical guide in R. STHDA. Kemp, A., 1995. The Hornbills: Bucerotidae. Oxford University Press, Oxford. 302 pp.

Lerm, R. E., Ehlers Smith, D. A., Thompson, D. I., & Downs, C. T. (2023). Human infrastructure, surface water and tree cover are important drivers of bird diversity across a Savanna protected area-mosaic landscape. Landscape Ecology, 38(8), 1991–2004. 10.1007/s10980-023-01674-2

Lloyd, T. J., Fuller, R. A., Oliver, J. L., Tulloch, A. I., Barnes, M., & Steven, R. (2020). Estimating the spatial coverage of citizen science for monitoring threatened species. Global Ecology and Conservation, 23, e01048. 10.1016/j.gecco.2020.e01048

Lok, A. F., Ang, W. F., Ng, B. Y. Q., Leong, T. M., Yeo, C. K., Tan, H. T. W. (2013). Native Fig Species As A Keystone Resource For The Singapore Urban Environment. Raffles Museum of Biodiversity.

Lüdecke D (2021)_sjPlot: data visualization for statistics in social science. R package version 2.8.10. https://CRAN.R-project.org/ package=sjPlot.

Lüdecke D (2023). sjPlot: Data Visualization for Statistics in Social Science. R package version 2.8.15, https://CRAN.R-project.org/package=sjPlot.

Marsden, S. J., & Pilgrim, J. D. (2002). Factors influencing the abundance of parrots and hornbills in pristine and disturbed forests on New Britain, PNG. Ibis, 145(1), 45–53. 10.1046/j.1474-919x.2003.00107.x

McCullagh, P. and Nelder, J. A.. 1989. *Generalized Linear Models*. Chapman Hall Miller, J. (2010). Species distribution modeling. *Geography Compass*, *4*(6), 490-509. 10.1111/j.1749-8198.2010.00351.x

Ng, S., & Cremades, M. (2023). The re-introduction of the oriental pied hornbill to Singapore. Peace with Nature, 267–273. 10.1142/9789811282027_0034

Ng, S., Lai, H., Cremades, M., Lim, M. T., Mohammed Tali, S. (2011). Breeding observations on the Oriental Pied Hornbill in nest cavities and in artificial nests in Singapore, with emphasis on infanticide-cannibalism. The Raffles Bulletin of Zoology Supplement, 24: 15–22.

Pearson, R. G., Raxworthy, C. J., Nakamura, M., & Townsend Peterson, A. (2006). Predicting species distributions from small numbers of occurrence records: a test case using cryptic geckos in Madagascar. Journal of Biogeography, 34(1), 102–117. 10.1111/j.1365-2699.2006.01594.x

Prudic, K., McFarland, K., Oliver, J., Hutchinson, R., Long, E., Kerr, J., & Larrivée, M. (2017). EButterfly: Leveraging massive online citizen science for butterfly conservation. Insects, 8(2), 53. 10.3390/insects8020053

R Development Core Team (2023) R: A Language and Environment for Statistical Computing. R Foundation for Statistical Computing, Vienna. https://www.R-project.org

SiReNT - Singapore satellite positioning reference network - Plane coordinate system - SVY21. (2020, June 8). https://app.sla.gov.sg/sirent/About/PlaneCoordinateSystem.

Spatial join (Analysis). (2023). https://pro.arcgis.com/en/pro-app/latest/tool-reference/analysis/spatial-join.htm

Strange, B. C. & O’Dempsey, T. (2022). A note on oriental pied hornbill reintroduction in Singapore and its dispersal from 2010-2021. Hornbill Natural History and Conservation. Vol. 3: 28–31,2022. https://iucnhornbills.org/wp-content/uploads/2022/07/HNHC_3_Strange-and-ODempsey.pdf

Sullivan, B. L., Wood, C. L., Iliff, M. J., Bonney, R. E., Fink, D., & Kelling, S. (2009). EBird: A citizen-based bird observation network in the biological sciences. Biological Conservation, 142(10), 2282–2292. 10.1016/j.biocon.2009.05.006

Tan, H. H. (2010). Spider-Feeding Behaviour Of The Oriental Pied Hornbill. Nature In Singapore. 3: 283–286

The Comprehensive R Archive Network. https://cran.r-project.org/web//packages/sjPlot/vignettes/plot_marginal_effects.html

Tiago, P., Pereira, H. M., & Capinha, C. (2017). Using citizen science data to estimate climatic niches and species distributions. Basic and Applied Ecology, 20, 75–85. 10.1016/j.baae.2017.04.001

Trisurat, Y., Chimchome, V., Pattanavibool, A., Jinamoy, S., Thongaree, S., Kanchanasakha, B., Simcharoen, S., Sribuarod, K., Mahannop, N., & Poonswad, P. (2013). An assessment of the distribution and conservation status of hornbill species in Thailand. Oryx, 47(3), 441–450. 10.1017/s0030605311001128

Urban Redevelopment Authority (URA). https://www.ura.gov.sg/-/media/Corporate/Planning/Master-Plan/MP19writtenstatement.pdf

Wei, T., Simko, V. (2021). R package ’corrplot’: Visualization of a Correlation Matrix. (Version 0.92), https://github.com/taiyun/corrplot.

Wei, T., & Simko, V. (2021). An introduction to corrplot package. The Comprehensive R Archive Network. https://cran.r-project.org/web/packages/corrplot/vignettes/corrplot-intro.html

Wong, T. W. (2011) The Singapore Hornbill Project: A Great But Simple Idea. The Raffles Bulletin Of Zoology. Supplement No. 24: 3–4

